# Resolving the full set of human polymorphic inversions and other complex variants from ultra-long read data

**DOI:** 10.1101/2025.05.27.656315

**Authors:** Ricardo Moreira-Pinhal, Konstantinos Karakostis, Illya Yakymenko, Oscar Conchillo, Maria Díaz-Ros, Andrés Santos, Miquel Àngel Senar, Jaime Martínez-Urtaza, Marta Puig, Mario Cáceres

## Abstract

Inversions are a unique type of balanced structural variants (SVs) with important consequences in multiple organisms. However, despite considerable effort, this and other complex SVs remain poorly characterized due to the presence of large repeats. New techniques are finally allowing us to identify the full spectrum of human inversions, but the number of individuals analyzed is still quite limited. Here, we take advantage of Oxford Nanopore Technologies (ONT) long reads to characterize an exhaustive catalogue of 612 candidate inversions between 197 bp and 4.4 Mb of length and flanked by <190-kb long inverted repeats (IRs). For that, we developed a bioinformatic package to identify inversion alleles reliably from long read data. Next, using a combination of different DNA extraction, library preparation, and ONT sequencing protocols, we showed that ultra-long reads (50-100 kb) and adaptive sampling are an efficient method to detect most human inversions. Lastly, by analyzing ONT data from 54 diverse individuals, 87-99% of the inversions could be genotyped in each sample, depending mainly on read and IR length and genome coverage. Both orientations were observed for 155 of the analyzed regions (frequency 0.01-0.49), which multiplies by three the polymorphic IR-mediated inversions studied in detail so far. Moreover, we found more than 300 additional independent SVs in the studied regions and resolved several complex rearrangements. Our work therefore provides an accurate benchmark of those inversions that typically escape most analyses, improving existing resources, such as the Pangenome. In addition, it demonstrates the potential of nanopore sequencing to determine the functional impact of missing human genomic variation.

## INTRODUCTION

Over the past few decades, there has been a global effort to identify all human genomic variation and its association with phenotypic traits and disease susceptibility in different populations (The 1000 Genomes Project Consortium 2015; Collins et al. 2020; Ebert et al. 2021; Karczewski et al. 2020; Choudhury et al. 2020; Choi et al. 2023). However, structural variants (SVs), which involve changes of more than 50 bp, are often generated through non-allelic homologous recombination (NAHR) between large repeats and tend to occur in complex regions harboring many segmental duplications (SDs), making their detection very difficult by most techniques (Alkan et al. 2011; Logsdon et al. 2020; Ebert et al. 2021). Thus, a big fraction of SVs are still under-studied and new high-throughput methods to determine reliably their genotypes in multiple individuals are needed.

This is especially problematic for inversions, which are generally undetected with most sequencing techniques due to their balanced nature, changing only the sequence order (usually with no gain or loss of DNA), and the frequent presence of highly identical inverted repeats (IRs) at their breakpoints (Puig et al. 2015; Ebert et al. 2021). In addition, compared to the rest of SVs, inversions are unique as they inhibit recombination between the two orientations, leading to potential negative consequences on fertility (Puig et al. 2015). Therefore, those that reach a certain frequency in natural populations could be more likely to have positive functional effects that compensate for these costs. In fact, a growing number of inversions have been involved in adaptation and phenotypic variability in diverse organisms, from plants to birds (Wellenreuther and Bernatchez 2018), including humans (Campoy et al. 2022). Moreover, around one third of human inversions are associated with gene expression or epigenetic changes, and they tend to have larger effects than other variants (Giner-Delgado et al. 2019; Puig et al. 2020; Lerga-Jaso et al. in prep.).

Large-scale projects using a combination of novel genomic techniques, such as long-read sequencing, Strand-Seq or Bionano optical maps (Levy-Sakin et al. 2019; Chaisson et al. 2019; Audano et al. 2019; Ebert et al. 2021; Porubsky et al. 2022) are finally making it possible to identify the full spectrum of human inversions. However, each technique has its own limitations and error sources. In addition, in most cases, just a reduced number of individuals has been studied, which hinders the analysis of the effects of the detected variants and their association with phenotypic traits. To overcome these problems, different methods have been developed to genotype accurately specific inversions, either individually (e.g. regular PCR, inverse PCR (iPCR) or droplet digital PCR (ddPCR)) or in sets of several of them together (e.g. multiplex ligation-dependent probe amplification (MLPA) or inverse MLPA (iMLPA)) (Giner-Delgado et al. 2019; Puig et al. 2020). This has allowed the detailed analysis of up to 134 inversions in 95-551 individuals from diverse human populations, including 54 NAHR inversions mediated by IRs ranging from 140 bp to 134 kb (Lerga-Jaso et al. in prep.). Nevertheless, the variants that can be studied with each method are extremely dependent on inversion and IR characteristics, such as their length or the presence of restriction enzyme target sites, which means that a single technique cannot detect all inversions and a significant fraction of them are still not characterized. In addition, many NAHR inversions have appeared recurrently in the human lineage and they are not linked to nearby SNPs or particular haplotypes (Giner-Delgado et al. 2019; Puig et al. 2020; Porubsky et al. 2022). Consequently, they cannot be imputed reliably using existing data sets and their potential effects have not been detected in common genome-wide association studies (GWAS).

Long-read sequencing methods, such as PacBio and Oxford Nanopore Technologies (ONT), represent a unique opportunity for the complete characterization of complex genomic variants (Logsdon et al. 2020; De Coster et al. 2021; Mastrorosa et al. 2023). Specifically, ONT sequencing is based on identifying the nucleotide bases as individual DNA molecules pass through multiple nanopores, which can generate extremely long reads of up to several Mb in a relatively short time (Logsdon et al. 2020). Despite its higher error rate, ONT ultra-long reads (>50-100 kb) have been instrumental in the generation of the new Telomere-to-Telomere (T2T) and Pangenome references, helping to resolve centromeres and other hard-to-characterize regions (Nurk et al. 2022; Liao et al. 2023). Furthermore, adaptive sampling promotes the sequencing of targeted molecules in real-time (Payne et al. 2021), enriching regions of interest in a simple and cost-effective way. This strategy has already been applied to identify rare genomic rearrangements and genotype difficult-to-study variants, such as tandem repeats (Miller et al. 2021; Stevanovski et al. 2022). Also, some studies are starting to generate long-read resources of multiple individuals of diverse populations and diseases (De Coster et al. 2021; Beyter et al. 2021; Wu et al. 2021; Gustafson et al. 2024; Schloissnig et al. 2024), although in most cases read length is moderate. Therefore, it is necessary to undertake further analyses to establish optimized protocols that can be widely adopted and define the current limits of these technologies. Also, it is important to develop easy to use bioinformatic tools that can extract accurate information of different types of genomic variants from these new data.

In this work, we take advantage of long read information to generate an accurate catalogue of human polymorphic inversions generated by NAHR spanning a wide range of IR lengths and to obtain an initial estimate of their frequency and distribution. Moreover, we investigate the applicability of ONT ultra long reads for efficient genotyping of inversions, as a first step to characterize their functional impact. Finally, we compare our results with other studies and the new Pangenome draft to check how well these variants are represented.

## RESULTS

### Generation of a complete inversion dataset

In order to build a complete dataset of IR-mediated human inversions, candidate inversion regions were selected from four different sources: (1) 58 validated inversions, of which 55 had already been genotyped in multiple individuals (Giner-Delgado et al. 2019; Puig et al. 2020; Lerga-Jaso et al. in prep.); (2) The merging of 1528 inversion predictions from five recent studies (Levy-Sakin et al. 2019; Chaisson et al. 2019; Audano et al. 2019; Ebert et al. 2021; Porubsky et al. 2022) that identify SVs using long-read sequencing, Strand-Seq or optical mapping in a total of 195 individuals; (3) Previous paired-end mapping (PEM) predictions that could not be analyzed with available methods or were identified as possible human genome assembly errors caused by highly-identical IRs (Korbel et al. 2007; Martínez-Fundichely et al. 2014; Vicente-Salvador et al. 2017); and (4) Pairs of inverted SDs annotated in the human reference genome (GRCh38/hg38) that have the potential to generate inversions by NAHR (see Methods). Predictions from the different sources were merged into independent potential inversions when their breakpoints were located within or close to the same pair of IRs. Also, we selected only inversion candidates with at least one IR copy smaller than 200 kb and not located in very complex regions full of repeats and SDs (which excludes most of the centromeric and telomeric sequences) or gaps in hg38, where specific probes could be designed around the breakpoints (see below).

Of the merged list of inversion predictions by recent studies, 624 regions were filtered out because they did not meet the above criteria (mostly because no IRs were identified in the region or sequence was extremely complex) and 11 more due to unexpected complex behavior during inversion genotyping. This resulted in 148 new possible independent inversions with IRs that could be tested (including 65 with breakpoints within or close to the same IR pair that were predicted by at least 3 independent studies), not counting 58 matching those already known (Lerga-Jaso et al. in prep.). In addition, there were 87 PEM inversion predictions overlapping IRs (Martínez-Fundichely et al. 2014) that were not present in the previous set. Finally, we selected 319 pairs of inverted SDs 1-50 kb long and distance of <250 kb in which no inversion predictions have been made so far. In total, 612 regions with potential inversions ranging from 197 bp to 4.4 Mb and flanked by IRs between 140 bp and 190 kb were included in the analysis (Supplemental Table S1), which constitutes the most exhaustive catalogue of candidate human NAHR inversions to date.

### Development of a new inversion genotyping package from long reads

Thanks to the sequence contiguity, ONT ultra-long reads offer the opportunity to analyze a wide range of inversions and resolve other challenging variants, without being affected by sequencing error rates or other methods limitations (Giner-Delgado et al. 2019; Puig et al. 2020). Thus, to extract this information easily and reliably, we have developed a novel inversion genotyping software package called GeONTIpe. The genotyping strategy is based on the detection of reads spanning inversion breakpoints and determining their sequence order and orientation by blasting specific probes at either side of each IR (A, B, C and D) (Figure 1A). That way, we can identify reads with A-B/C-D combinations that support the reference orientation (O1) or A-C/B-D combinations that support the inverted orientation (O2) (Figure 1B).

**Figure 1.**
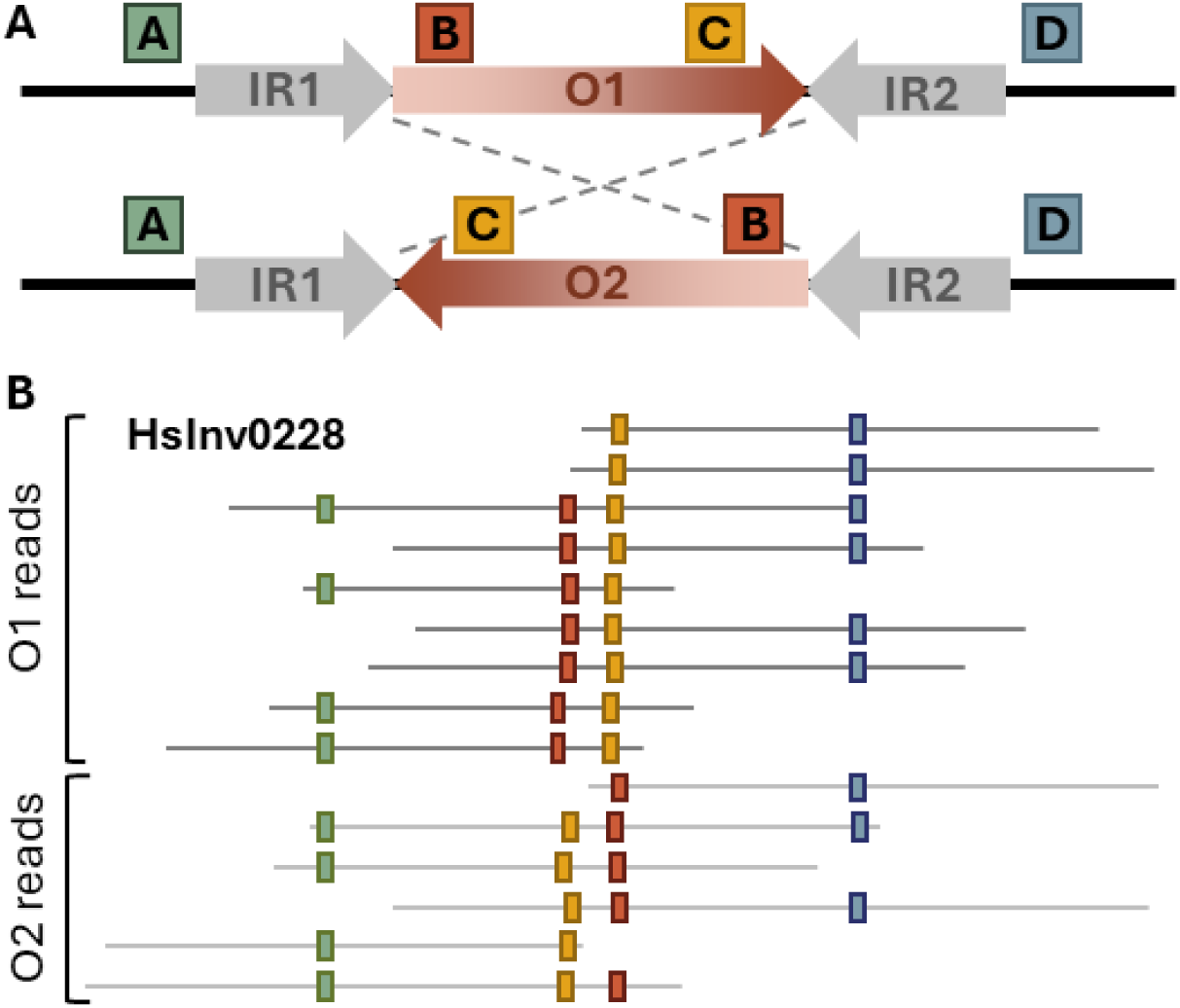
Inversion genotyping based on ONT long reads. **A.** Schematic representation of candidate inversion regions analyzed in this work, with the inverted region shown as a red gradient flanked by two inverted repeats (IR) in grey. Probes located at both sides of each IR are indicated as colored rectangles and have different order in O1 and O2 orientations. **B.** Example of HsInv0228 inversion genotyping results in an heterozygote sample, showing ONT individual reads as dark grey (O1) or light grey (O2) lines and the included probes indicated with colored rectangles. Probe order together with read number and proportion of reads supporting each allele determines the inversion genotype.

The analysis has several steps: quality filtering of the reads, mapping against hg38, generation of inversion genotypes, and identification of additional SVs (see Methods; Supplemental Figure S1). One of the key steps is the design of the probes to identify unique sequences flanking the breakpoint IRs, which were manually revised and updated to avoid specificity problems and possible errors caused by the presence of unexpected haplotypes. This allowed us to interrogate accurately 612 potential inversions, including 40 inversions analyzed only using one breakpoint due mainly to the presence of large IRs or the lack of specific probes. Once the probes were mapped to the reads, inversion genotype was determined based on the number of reads supporting each orientation as homozygote, heterozygote or low confident (see Methods). Moreover, to recover additional genotypes, in cases where possible homozygotes were supported just by 2-5 reads, known SNPs in the region were analyzed to check for the presence of two alleles. Finally, we took advantage of the probe distance information to identify additional insertions (increased distance) or deletions (reduced distance) in the analyzed regions. Therefore, the developed method makes it possible to resolve a large majority of human inversions mediated by a wide range of IRs for the first time, while it characterizes also additional SVs in these regions.

### Benchmark of ONT sequencing strategies for inversion genotyping

In parallel, to assess the most efficient methodology for genotyping a complete set of human inversions on a single experiment, we tested several ONT sequencing protocols that generate ultra-long reads using a MinION device. The analysis was multi-dimensional and includes two main sets of experiments on four human female samples of diverse origins for which most of the validated inversions had been previously analyzed (Lerga-Jaso et al. in prep.): NA12156 (European), NA18505 and NA18508 (African), and NA18956 (East Asian) (Figure 2A). First, we used the Monarch® HMW DNA extraction method (NEB) to compare the genotyping efficiency of whole-genome sequencing (WGS) and adaptive sampling (AS) of two sets of inversions: an initial set, including 132 candidate inversions plus 40-kb flanking regions (AS2), and an extended one, targeting additional predicted inversions and inverted SDs plus 70 kb flanking regions (AS3), which covers almost all the inversions interrogated in the genotyping pipeline (Supplemental Table S1). Second, we evaluated the effect on read length and total output of an alternative phenol-based genomic DNA purification protocol using AS3.

**Figure 2.**
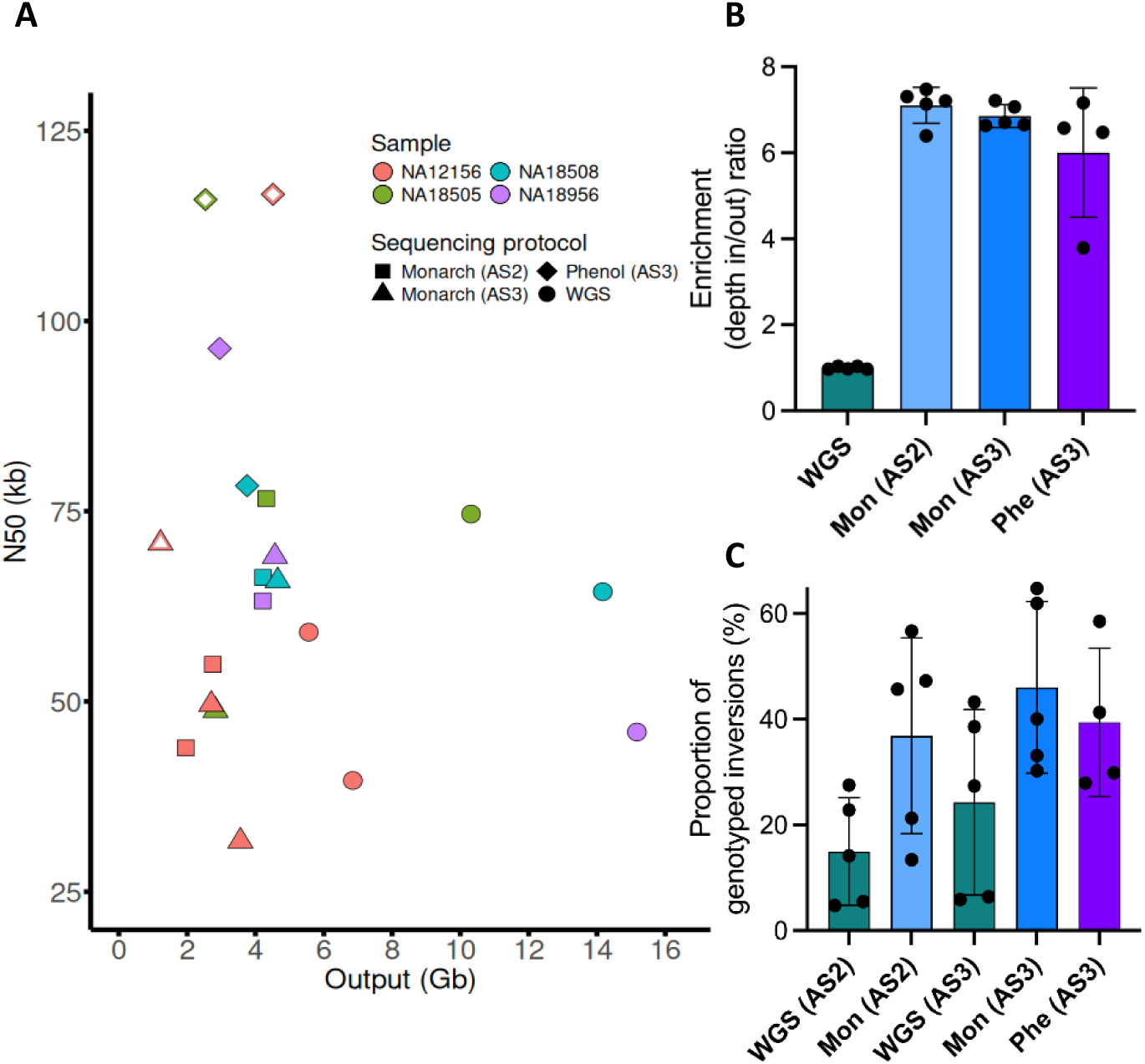
Summary of ONT sequencing experiment results. **A.** Comparison of the N50 value and the total output of each sequencing run of the four analyzed samples (in different colors) (Supplemental Table S2). Diverse sequencing protocols using whole genome sequencing (WGS) or adaptive sampling (AS2/AS3) and DNA isolation methods (Monarch HMW or phenol) are indicated by different shapes, whereas empty symbols indicate the use of fresh cells as starting material. **B.** Enrichment ratio between the read depth inside and outside the targeted regions (depth in/out) of each sample for diverse types of sequencing (black dots), showing a 6-7-fold average enrichment in AS2 or AS3 compared to WGS (colored bars). **C.** Average proportion of genotypes obtained for the analyzed inversion regions normalized to the number of AS2 or AS3 targets (%, colored bars). Values for each sample (black dots) show considerable variation due mainly to sequencing output differences.

In the different sequencing experiments with the Monarch extracted DNA, we obtained an N50 of 32-79 kb, with consistent values when the same library was sequenced and no significant differences between WGS and AS (e.g., WGS and AS2 in samples NA12156 and NA18505 or AS2 and AS3 in samples NA18508 and NA18956) (Figure 2A; Supplemental Table S2). However, the phenol-based DNA extraction method resulted in significantly larger N50, ranging from 79 to 118 kb (Figure 2A). Also, long-term storage of the cells appears to reduce genomic DNA integrity and read length, with DNA isolated from fresh cells by phenol resulting in higher N50 values than DNA isolated from cells stored at −80 °C for a year (mean of 117.7 vs 87.9 kb, respectively), whereas the N50 of the DNA extracted with the Monarch kit from NA12156 cells kept for 8 years at −80 °C showed clearly the lowest N50 (50.6 kb on average) (Figure 2A). In terms of output, both Monarch AS2 and AS3 experiments yielded an average of 3.3 and 3.1 Gb compared to 9.5 Gb for WGS, which corresponds to a ratio of 2.4, 2.0 and 5.5 Mb/nanopore, respectively (Figure 2A; Supplemental Table S2). This drastic effect on output could be caused from reduced activity and lifetime of nanopores during AS. Thus, our results suggest that read length depends mainly on sample preparation, while the amount of sequence data is reduced by ∼3-fold in AS compared to WGS.

Regarding the efficiency and specificity of AS sequencing, we observed a similar average increase on the coverage of the targeted inversion regions for the Monarch AS2 (7.1-fold) and AS3 (6.9-fold) results, which is quite consistent across samples (Figure 2B). Dividing the genome into 10-kb windows showed a clearly different distribution of the average fold enrichment between targeted and non-targeted windows (Supplemental Figure S2; Supplemental Figure S3A) and a significant correlation of the mean enrichment of the different target regions common in AS2 and AS3 (*r* = 0.35, *P* < 0.001; Supplemental Figure S3B). Still, in both AS datasets a small proportion of non-targeted windows display increased coverage values comparable to that of targeted regions (Supplemental Figure S2; Supplemental Figure S3A). To address this, we analyzed the presence of different types of repeat sequences that could be driving off-target effects in the top 1% non-targeted windows with highest coverage in AS2 and AS3. Indeed, 71.2% and 85.5% of off-target windows in AS2 and AS3 showed a significant enrichment of repetitive elements, SDs or both (Supplemental Figure S3C), with more AS3 off-target windows enriched by SDs compared to AS2 (74.5% vs. 45.2%), probably due to the higher SD content of the AS3 targeted regions. Next, we searched for sequence homology between the AS3 targets and the top 1% non-targeted windows. From a total of 2738 AS3 off-target windows, 1399 (51.1%) had at least one highly identical Blast hit with the targeted sequences (75.46%). Thus, a high proportion of AS off-target effects could be due to homology with repetitive elements or other sequences.

When considering inversion genotyping, Monarch AS2 and AS3 sequencing experiments provide respectively an average of 1.9 and 1.7 times more informative reads for inversion genotyping than WGS (Supplemental Table S2), which results from a trade-off between the higher enrichment and the lower output of AS. Similarly, depending on each run results, up to 65% of the targeted inversions could be successfully genotyped in a single MinION flow cell, and on average this value is 1.9-2.5-times higher in AS2 (mean = 36.9%) or AS3 (mean = 46.0%) than in WGS (mean = 15.0% and 24.3% for AS2 and AS3 targets, respectively) (Figure 2C; Supplemental Table S2). Even though phenol-extracted DNA generated significantly longer reads, with N50 >100 kb from fresh cells, this did not result in better inversion genotyping (mean = 39.4%), probably because the longer molecules yielded less total sequence reads (Figure 1A and 1C). When the data of the different sequencing experiments of the same sample are merged, 87-91% of the inversions could be genotyped in each individual.

### Large-scale inversion genotyping in humans

To obtain a good picture of human NAHR polymorphic inversions, we applied the inversion genotyping pipeline to a diverse set of 54 individuals with publicly available ONT data, including those sequenced in this study (Supplemental Table S3). The analysis showed a high genotyping success (>87%) of the candidate inversion regions in all the samples (Figure 3A). As expected, the sample genotyping rate is correlated with the sequencing depth and especially N50, with 99% of the inversions being genotyped in the samples with N50 >100 kb (Figure 3B). Similarly, the distance between probes at both sides of the breakpoints reduces inversion genotyping success due to a lower number of reads crossing the IRs (Figure 3C). In fact, inversions with probes separated by >100 kb could be genotyped on average in only 47.0% of the samples.

**Figure 3.**
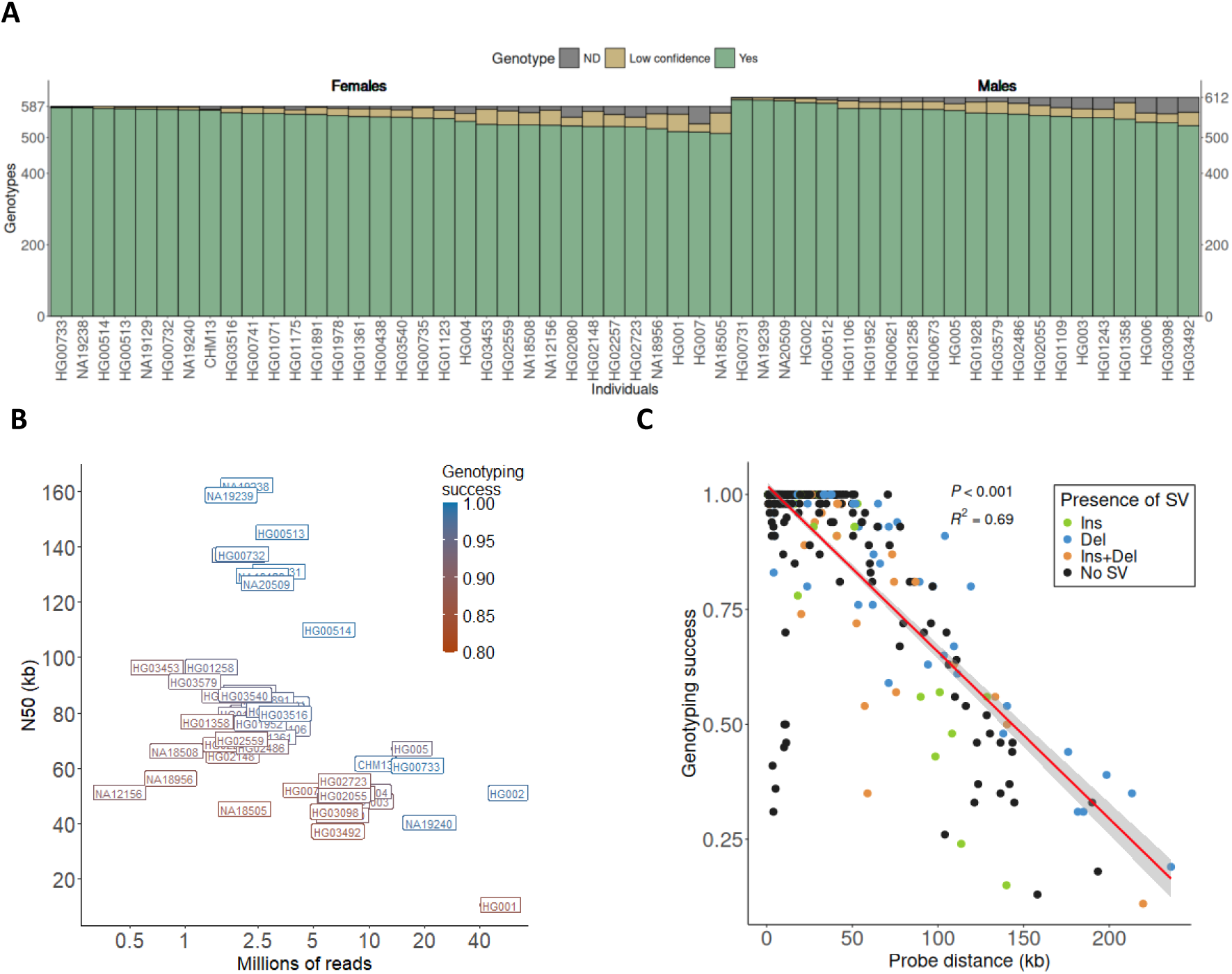
Genotyping success of candidate inversion regions from ONT data in 54 individuals. **A.** Summary of inversion genotyping results by individual, with complete genotypes in green, low confident genotypes due to a reduced number of reads in yellow, and those with no supporting reads (not determined, ND) in grey. The maximum number of genotypes is 612 for males and 587 for females, excluding 25 predicted inversions in Chr. Y. **B.** N50 values and total number of reads for each sample, showing the genotyping success rate in a red to blue scale. **C.** Genotyping success of candidate inversion regions as a function of the average distance between the probes flanking the analyzed breakpoints. Additional SVs detected in the regions between the probes, some of which can affect genotyping success, are indicated in different colors.

We detected the two orientations in at least one of the 54 individuals (not counting the hg38 reference genome to avoid possible assembly errors) in 155 of the 612 inversion regions tested (25.3%). The minor allele frequency (MAF) of the detected inversions ranged from 0.01 to 0.49, with 78.1% of the polymorphic inversions being widespread across populations (Supplemental Table S1; Supplemental Table S3). However, the fraction of regions identified as polymorphic inversions varied depending on the source of the predictions, with two alleles detected in this set of individuals in all but three of the 58 previously known inversions (94.8%), 54.7% of the additional inversion predictions merged from five different studies, 13.8% of other PEM predictions that had not been validated before, and, as expected, only 2.2% of the inverted SD pairs (Figure 4A). The previously known inversions not found here correspond to two very low-frequency inversions (HsInv0486 and HsInv0626), and HsInv1869, which could be an hg38 assembly error since only inverted alleles are found in all analyzed samples (Supplemental Table S3). Regarding the predictions from recent SV studies, a similar proportion of real inversions was found in those using a multiplatform approach or long reads (74.8-80.8%), with a slightly lower success for the study relying only in Bionano optical maps (66.2%), although it is also the one analyzing a larger number of samples (Figure 4B). The remaining 67 predictions not detected as polymorphic inversions were predominantly predicted only in one or two of these studies (47), which could represent false predictions or low frequency variants. Those found in three or more studies correspond mainly to possible hg38 reference genome assembly errors, in most of which all the individuals genotyped have the O2 orientation, including 13 that had been already confirmed experimentally (Vicente-Salvador et al. 2017; Antonacci et al. 2010). Also, there were a few inversion candidates in complex regions with different breakpoint SD pairs that resulted in different predictions in which only one was validated, or that showed low genotyping success due to the length of the repeat blocks. Interestingly, by checking additional individuals using ONT data, two previously validated reference genome errors (Vicente-Salvador et al. 2017) turned out to be low frequency inversions (HsInv0057, HsInv1119). Finally, the ONT data analysis confirmed that many previous PEM predictions were probably errors caused by variation within SDs among individuals and complex regions (Figure 4A).

**Figure 4.**
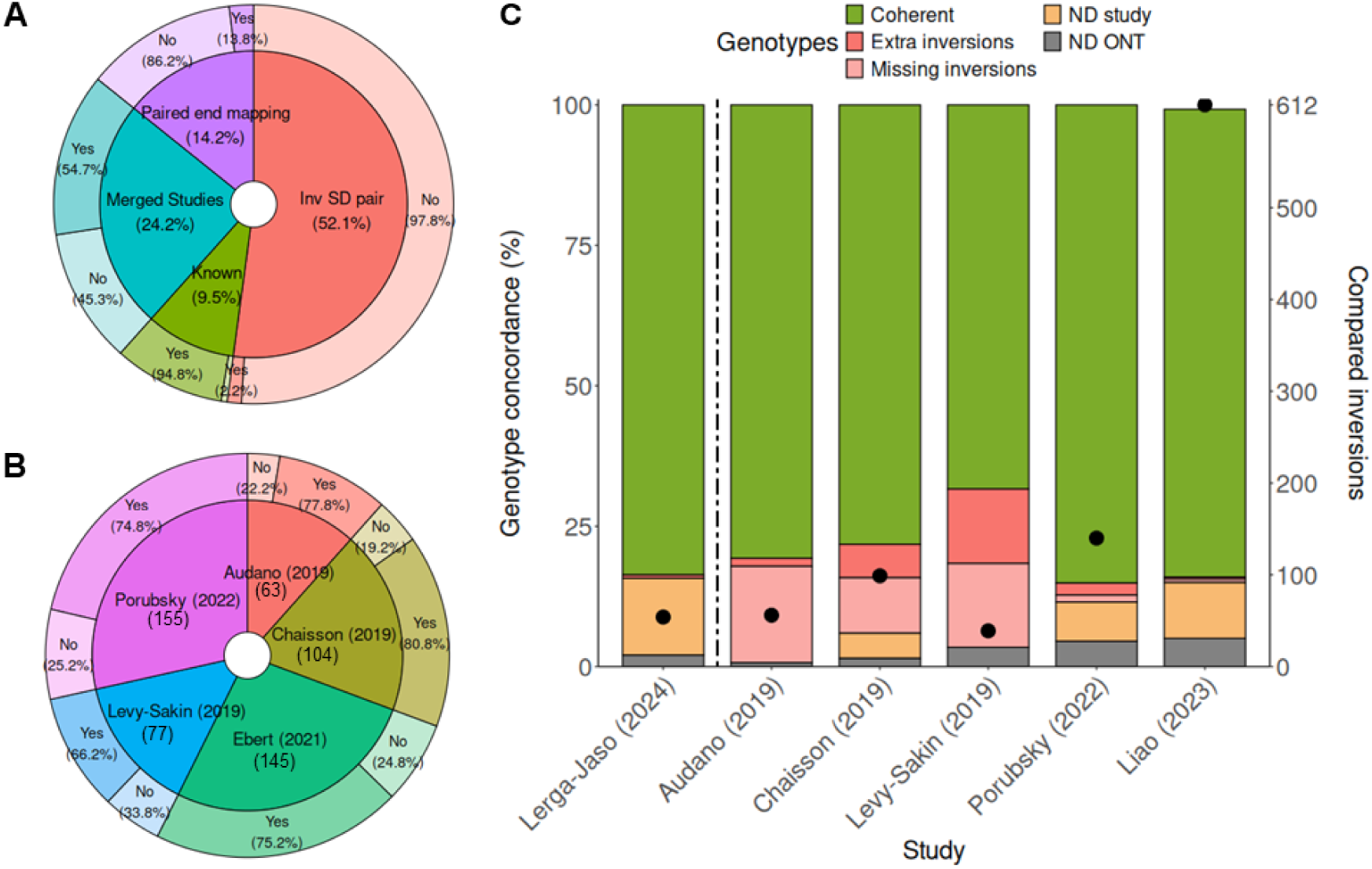
Comparative analysis of inversion genotypes obtained in this and previous studies. **A.** Pie chart representing the proportion of the 612 analyzed regions derived from different sources (inner circle, see Supplemental Table S1) and those that were detected as polymorphic inversions or not in our dataset (outer circle)**. B.** Pie chart representing the total number of candidate regions from each of the merged SV studies included in our analysis (inner circle) and the percentage detected as polymorphic inversions or not (outer circle). **C.** Concordance of ONT genotypes with PCR-based experimental data (separated by dashed line) and five other available inversion prediction studies analyzing the same samples and inversion regions (left Y axis). Genotypes are classified as concordant, having extra or missing inverted alleles, or not determined (ND) with ONT data or in the other study used for comparison. Black dots represent the number of inversions compared in each dataset (right Y axis).

### Accuracy of ONT inversion genotyping

We performed different comparisons to demonstrate the accuracy of the methodology. First, we checked the genotypes of the polymorphic inversions in five parent-child trios included in our dataset (Supplemental Table S3). Only 2 of the 652 trio comparisons showed discordant results (99.7% concordance): HsInv0808 in the HG002 Ashkenazi trio child that showed a single O2 read and 31 O1 reads, which was insufficient to call it as heterozygote, and HsInv0278 in the HG00731 Puerto-Rican trio father, with 35 O2 reads, that in previous studies already showed discrepant calls as O2 homozygote (Chaisson et al. 2019) and as heterozygote (Porubsky et al. 2022). In addition, we matched 54 inversions that have been experimentally genotyped by different PCR-based methods in 10 individuals included in the present analysis (Giner-Delgado et al. 2019; Puig et al. 2020; Lerga-Jaso et al. in prep.). In total, 452 of the 455 genotypes compared (99.3%) were concordant (Figure 4C), and all three discrepancies correspond apparently to errors in the experimental data. These involved a new inverted duplication identified from the long read data instead of the expected inversion (HsInv0397 in NA12156), a complex duplicated region (HsInv0348 in NA18508) or an inversion with a high genotyping error rate (Giner-Delgado et al. 2019) in which the 1000 Genome Project (1KGP) high-coverage data supports also the ONT results (Byrska-Bishop et al. 2022) (HsInv0045 in NA20509). Therefore, ONT inversion genotyping is able to correct errors of other methods, being even more reliable than some experimental genotypes.

Next, we compared our results with those of four previous studies, taking into account the full genotype, if available, or just the presence or absence of the inverted allele (Levy-Sakin et al. 2019; Chaisson et al. 2019; Audano et al. 2019; Porubsky et al. 2022) (Figure 4C). A lower genotype concordance was observed with older studies, that were based either in one detection technique, such as Bionano optical maps (70.8% from 39 inversion predictions in 6 individuals) (Levy-Sakin et al. 2019) and PacBio long reads (81.3% from 56 inversion predictions in 5 individuals) (Audano et al. 2019), or a multi-platform approach integrating the previous two techniques and Strand-Seq (83.2% from 99 inversion predictions in 9 individuals) (Chaisson et al. 2019). Conversely, a recent large multi-platform analysis using the same techniques shows 96.2% genotype concordance from 140 inversion predictions in 12 individuals (Porubsky et al. 2022). Nevertheless, one fourth (25.2%) of the 155 polymorphic inversions found were missed in that study.

Finally, we checked how well the complete set of inversions are represented in the available diploid genomes of 14 males and 19 females from the new Pangenome draft (Liao et al. 2023). We could compare 84.2% (16614) of all the genotypes, which showed almost perfect concordance with our results (98.8%, Figure 4C). However, most of the remaining genotypes could not be compared mainly due to the absence of assembled sequences crossing the breakpoints in some inversions and individuals in the Pangenome (10.7%, 2120). Thus, even though the Pangenome provides a good representation of polymorphic inversions, it still misses information in a significant fraction of regions or individuals.

### Identification of other SVs and characterization of complex regions

After mapping the probes on each read, the comparison of the actual distances between them with those expected in the hg38 reference genome or its inverted version allowed us to identify additional SVs in the tested regions. In total, we detected 577 deletions and insertions of different sizes (between 250 bp and 469 kb) in either O1 or O2 chromosomes (Supplemental Table S4), which correspond to ∼322 independent SVs: 195 deletions, 89 insertions, and 38 possible CNVs (see Methods for details). Of those, there were 243 SVs between the probes flanking one breakpoint, 65 between the two internal probes within the inverted sequence, which could be only analyzed for part of the inversions shorter than 50-100 kb, and 14 deletions that remove one or more of the internal probes. Interestingly, regions with polymorphic inversions contain more additional SVs (137 SVs in 76 regions (49.0%), with an average of 1.8 SVs per region) than those without inversions (185 SVs in 140 regions (30.6%), with an average of 1.3 SVs per region). This suggests that rearrangements tend to accumulate in certain parts of the genome. In addition, although our data only analyzed presence or absence of each SV in the reads of an individual, their frequency ranges from those found in only 1 out of 54 analyzed individuals to those found in all of them, corresponding probably to an error or low-frequency haplotype in the reference genome (Supplemental Table S4), with 65.5% of them (211 SVs) detected in several individuals, which indicates that they are relatively common and that they are not false positives.

For some regions that accumulate several SVs, we combined the pipeline output and manual read analysis to identify multiple structural haplotypes. For example, in HsInv0401 there is an 8.7-kb polymorphic inversion and a 12.7-kb copy number variant (CNV) at one or both breakpoints that can have 2-5 copies per chromosome and result in nine configurations (Figure 5A). We also characterized other types of complex regions, like that of HsInv0822-0816-0384, where three nested polymorphic inversions can generate up to eight different haplotypes (Porubsky et al. 2022) of which we detected six. Similarly, in the HsInv0012-0659 region a polymorphic inversion, a CNV with 0-2 copies, two insertions, and a deletion generate at least five different haplotypes (Figure 5B). Thus, ONT data are capable of resolving precisely complex regions including different combinations of SVs.

**Figure 5.**
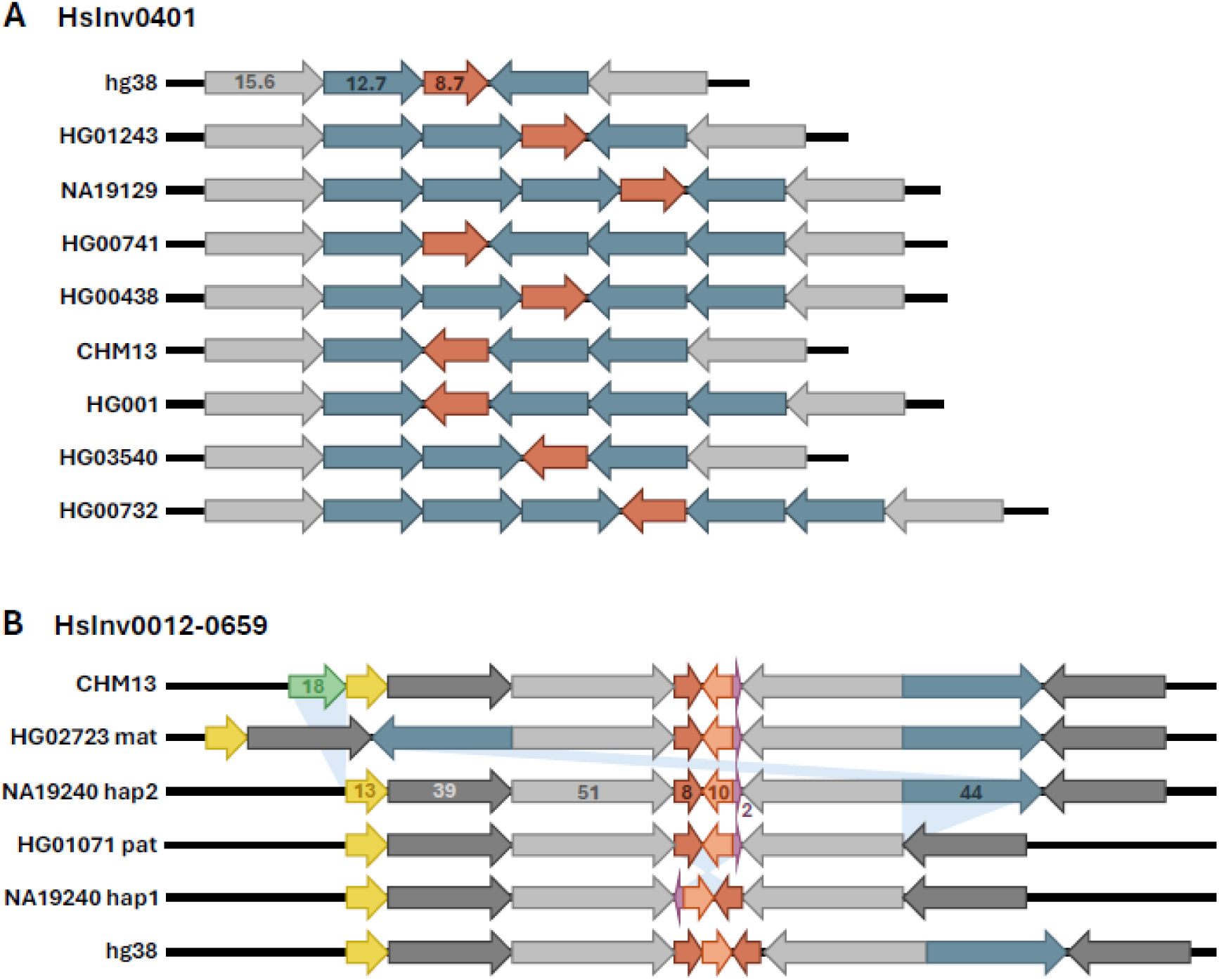
Examples of structural diversity at complex inversion regions resolved with ONT long reads. **A.** HsInv0401 includes a 12.7-kb CNV with up to five copies per chromosome (blue arrows) located within the inverted repeats at both sides of the inverted sequence (red arrow). **B.** HsInv0012-0659 includes a polymorphic inversion (red, orange and purple arrows), a 44-kb duplication and deletion (blue arrows) and an 18-kb insertion (green arrow). In this case, the hg38 reference genome sequence is not supported by any other assembly we analyzed so far and might represent an assembly error. In both images, grey arrows correspond to non-variable regions of inverted SDs and numbers indicate the size of the different segments in kb.

## DISCUSSION

During the last years considerable efforts have been dedicated to the identification of all genetic variation in humans (Sudmant et al. 2015; Byrska-Bishop et al. 2022; Levy-Sakin et al. 2019; Chaisson et al. 2019; Audano et al. 2019; Collins et al. 2020; Ebert et al. 2021; Porubsky et al. 2022; Gustafson et al. 2024; Schloissnig et al. 2024). However, a detailed characterization of the different types of SVs found is needed to determine precisely how many variants there really are and their mutational effects. Here, by doing a comprehensive analysis of the latest human SV information, we have created an accurate catalogue covering the large majority of human inversions mediated by IRs, which typically escape most analysis. In addition, we have developed a complete bioinformatic and experimental approach to genotype complex inversions not equaled by other available techniques. This includes a new easy-to-use bioinformatic package that generates highly reliable inversion genotypes from different types of long read data (Schloissnig et al. 2024). Furthermore, by testing different DNA extraction, library preparation and sequencing protocols, we have shown that the combination of ultra long ONT reads and AS constitutes an efficient method to analyze most human NAHR inversions in a single experiment, which reduces enormously the amount of time and effort required. Thus, taking advantage of ONT long read data from multiple individuals, we have been able to: (1) validate 155 polymorphic inversions, including several not previously detected; (2) identify more than 300 additional SVs in the analyzed regions; and (3) resolve complex regions with different rearrangements generating structurally diverse haplotypes that deserve further study.

In fact, ONT long reads represent a more direct way to identify inversions and characterize complex regions than other available techniques, since it is just based on the mapping of long stretches of DNA, which is more reliable than that of shorter sequences, especially for repeats. For example, around half of the IR-mediated inversions predicted in recent studies were not detected in our analysis (Figure 4A). Part of the discrepancies can be due to not interrogating the same individuals and others correspond to potential assembly errors in the human reference genome (Vicente-Salvador et al. 2017). However, our results suggest that there is still a significant proportion of false predictions, which is also illustrated by the 4-53 extra inversion calls in the few individuals compared in the different studies with respect to the ONT data (Figure 4C). This is especially evident in the case of older PEM inversion predictions (Martínez-Fundichely et al. 2014), with 86.2% of those not already validated or present in the recent studies not detected in our analysis (Figure 4C), despite half of the analyzed individuals in both cases are the same. Moreover, most previous studies capture only a fraction of the actual polymorphic inversions. Therefore, our high-quality inversion benchmark constitutes a very useful resource for SV detection projects and future attempts to represent completely human structural diversity (Wagner et al. 2022), such as the new human pangenome (Liao et al. 2023).

Another important contribution of this work is the generation of reliable genotypes for most inversions mediated by IRs, including many challenging variants that could not be analyzed before. Specifically, we show that ONT inversion results are even more accurate than experimental genotypes, allowing us to correct three existing errors from PCR based techniques (Giner-Delgado et al. 2019; Lerga-Jaso et al. in prep.). Long read mapping has the advantage that avoids some problems of other methods, like the need for restriction sites in specific positions, and it is more robust to different types of variation that cause genotyping errors (Giner-Delgado et al. 2019). Thus, the design of the assays is very flexible and can cover a wide range of inversions, with the only limitation being having reads longer than the IRs and finding uniquely-mapping probe sequences at each side. In addition, the actual sequence provides extra information that allows us to identify SNPs or resolve other SVs, as in the low frequency inverted duplication of HsInv0397 and other complex regions analyzed here. Regarding previous studies, apart from the Pangenome draft, we found the best genotype concordance with the most recent multiplatform analysis, which includes long reads (Porubsky et al. 2022). However, ONT sequencing achieves this performance with a lower cost and effort than the different techniques used.

Regarding the sequencing protocol optimization, we have shown that ONT AS strategy does obtain a significant and quite-specific enrichment of sequence reads from the regions of interest, which is somehow compensated by a lower total sequencing output of each run (probably due to reduced pore life) (Payne et al. 2021; Kovaka et al. 2021), resulting finally in just a ∼2-fold increase in the inversions genotyped compared to sequencing the whole genome. The combination of AS and the appropriate DNA extraction and library preparation methods to generate ultra long reads with N50 up to 70-100 kb allows us to identify consistently 40-60% of inversions in each individual MinION flow cell, which increases to ∼90% by pooling the data from 3-4 flow cells together. This is equivalent to the results obtained from other available high coverage genomes (Figure 3A) and it means that the whole set of inversions could likely be analyzed in a single PromethION flow cell.

As seen in Figure 3C, one of the main limitations is the size of inversion breakpoint IRs, with genotyping success decreasing quickly for IRs over 50 kb. For example, for most inversions with the largest IRs (>150 kb), less than one fourth of genotypes are obtained, unless there are deletions that reduce the distance between probes. Similarly, genotyping success of the samples increases as the reads get longer (higher N50), although good genotyping can also be obtained with lower N50 and more coverage, since there are still enough long reads to genotype most inversions (Figure 3B). In addition, according to our results, using fresh cells is important to extract the longest DNA molecules and increase N50 (Figure 1A). However, there is a well-known trade-off between read length and sequencing output with ONT, so that higher N50 values do not necessarily mean more genotyped inversions, as happens for the samples extracted using the phenol protocol (Figure 3C). Thus, ONT inversion genotyping depends on a balance between several factors with opposing effects, such as read length, number of sequences and sequence selection affecting pore survival.

Accurate genotypes of SVs in multiple individuals are crucial to determine their functional effects and their association with phenotypic traits and disease susceptibility. By extending the current work to the increasing amount of available ONT genome data (Gustafson et al. 2024; Schloissnig et al. 2024), future analyses will allow us to investigate a diverse set of inversions in a large number of individuals. These genotypes will help to improve inversion imputation in functional and phenotypic data, especially for recurrent inversions, for which there is very limited information (Giner-Delgado et al. 2019; Puig et al. 2020; Lerga-Jaso et al. in prep.; Porubsky et al. 2022), making possible to determine their functional effects more precisely. Also, the additional genotypes will provide a better estimate of inversion frequency and distribution across human populations. It is important to mention that there is still a small fraction of inversions with extremely long IRs (>150-200 kb) that cannot be detected with long reads and require more specialized techniques like Strand-Seq (Sanders et al. 2016; Porubsky et al. 2022). Nevertheless, ONT sequencing improves extraordinarily current inversion analysis and opens the door to the full characterization of one elusive type of human genetic variant for the first time. Finally, due to the potential of the technology, it is expected that continued advances and standardization of the methodology would likely improve sequence accuracy, output and targeting. This will likely result in reduced costs and increase its widespread use to analyze inversions and other complex variants in basic and translational research (Dixon et al. 2023; Xu et al. 2023; Miller et al. 2021; Stevanovski et al. 2022) contributing to a more complete picture of human genetic variation.

## METHODS

### Human samples

Lymphoblastoid cell lines (LCLs) of four unrelated females included in the 1KGP were obtained from the Coriell Cell Repository and they were cultured in T75 flasks at 37°C, 5% CO_2_ in DMEM medium, supplemented with antibiotics, 2 mM L-glutamine (Gibco/Invitrogen) and 10% fetal serum (HyClone), until saturation. Cells were harvested and pellets were stored at −80°C or used directly for DNA isolation. Also, we used available ONT sequencing data from 50 additional individuals of diverse origins generated mainly by the T2T Consortium (1) (Nurk et al. 2022), the Genome in a Bottle Consortium (7) (Zook et al. 2016), the Human Structural Variation Consortium (11) (Porubsky et al. 2022) and the Human Pangenome Reference Consortium (31) (Liao et al. 2023)) (Supplemental Table S1). All procedures that involved the use of human samples were approved by the Animal and Human Experimentation Ethics Committee (CEEAH) of the Universitat Autònoma de Barcelona.

### Generation of NAHR inversion dataset

To generate the dataset of candidate inversions mediated by NAHR, first we merged 58 previously characterized inversions with IRs (Giner-Delgado et al. 2019; Puig et al. 2020; Lerga-Jaso et al. in prep.) and 1528 inversion predictions from five different studies: Levy-Sakin et al. (2019) (314), Chaisson et al. (2019) (320), Audano et al. (2019) (220), Ebert et al. (2021) (315), and Porubsky et al. (2022) (359). IRs at the prediction breakpoints were identified from the UCSC Genome Browser hg38 human assembly SD track (https://genome.ucsc.edu) or by BLAST (>90% identity) of the extended inverted region against itself. Those predictions with both breakpoints located respectively within the same IR pair were considered to represent the same inversion. When there were several possible SD pairs overlapping the breakpoints, then different predictions were generated and tested. Second, we took all the inversion predictions generated by PEM of longer fragments (Korbel et al. 2007; Kidd et al. 2008) in the InvFEST database (Martínez-Fundichely et al. 2014) and removed those matching regions already identified in the previous step. The remaining predictions with inverted SDs or IRs at the breakpoints, assessed as before, were added to the dataset. In both cases, regions without IRs, containing complex repeat structures or large gene families, breakpoints within SD blocks >200 kb, or inverted regions <50 bp, were excluded from the analysis. Third, other inverted SDs with 90% identity that could result in inversions were obtained from the UCSC hg38 SD track, after filtering repeats longer than 50 kb or further apart than 250 kb from each other. Inverted SDs that had a reciprocal overlap larger than 80% were merged with BEDTools v2.30.0 and the coordinates of the merger were used. Only those regions not already included were added to the final NAHR inversion list. Finally, additional extra regions previously identified as assembly errors in the reference genome that had IRs at both breakpoints (Vicente-Salvador et al. 2017) were also included in the analysis.

### DNA isolation and library preparation

High molecular weight genomic DNA was obtained using two different methods by following the protocol specifications: (1) the Monarch HMW DNA Extraction Kit for Cells & Blood (New England Biolabs) starting from ∼6 million cells; and (2) a phenol-based isolation protocol optimized to obtain ultra-long reads for ONT sequencing (Gong et al. 2019) starting from 20-30 million cells. DNA was resuspended in 200 to 760 μl of EEB buffer (10 mM Tris-HCl, 1 mM EDTA pH 9 and 0.5% Triton X-100) provided in the library construction kit (see below). DNA concentration was measured shearing 10 μl by vortexing and using the Qubit Fluorometric Quantification and the AccuGreen Broad Range dsDNA Quantitation Kit reagents (Biotium). Sequencing genomic libraries were prepared using the Ultra-Long DNA Sequencing Kit V14 (ONT) following the manufacturer’s instructions. DNA libraries were dissolved in 300 μl of Elution Buffer (10 mM Tris-HCl, pH 8.0) provided by the kit and kept at 4 °C. During these steps, DNA samples were always handled with care using wide-bore tips, without being vortexed, shaken, or centrifuged at high speed to avoid DNA shearing.

### ONT sequencing

Sequencing was performed using a MinION Mk1B or Mk1C device and R10.4.1 flow cells (ONT). Flow cells ran approximately for 3-5 days with flushes every 24 hours and each time 37 μl of library were loaded into the flow cell. The MinKNOW software (version 22.12.5 to 23.04.3) was used to run the sequencing assays, and high-accuracy (HAC) basecalling was performed with Guppy (version 6.4.6 to 6.5.7). Real time adaptive sampling was done using MinKNOW in a Windows 10 2009 computer with an AMD Phenom II X4 955 processor, 16 GB of RAM and a PNY Nvidia GTX 1080 GPU. Two different versions of a FASTA file with the target sequences were used: AS2, an initial set including only 132 candidate inversions regions; and AS3, the final set that contains 587 of the 612 inversions interrogated (Supplemental Table S1). Targeted regions in AS2 and AS3 files spanned, respectively, 40 or 70 kb of sequence at both sides of the IRs at inversion breakpoints, excluding the actual IRs to avoid generation of non-informative reads that do not cross the breakpoint. Supplemental Table S6 summarizes the information of all the sequencing runs generated in this work, including tests using additional DNA extraction and library preparation methods for the NA18505 and NA18508 samples. All the available data were merged together in the final genotyping of these samples to maximize the number of inversions resolved.

### Inversion genotyping pipeline

As explained above, inversions were identified by mapping a set of four specific probe sequences flanking the inversion breakpoints on ONT reads (Figure 1A). Probes were designed by a combination of manual analysis and a custom script that generates 500-bp sequences with a 250-bp overlap along the inversion region (extending up to 100 kb at each side of the external limits of the breakpoints). Manual and automatic sequences were used as query on Blastn v.2.12.0 (Altschul et al. 1990) against the inversion region ± 500 kb in the hg38 and T2T reference genomes with parameters -perc_identity 80 and -qcov_hsp_perc 90, and those closest to the breakpoint and showing a single hit were selected as probes. Selected probes were tested for inversion genotyping (see below) and they were manually adjusted when needed to avoid specificity problems caused by unexpected variation.

The inversion genotyping pipeline consisted of multiple steps that use different computer programs and custom scripts. These programs and scripts, including the probe design, have been joined together in a new package named GeONTIpe (v1.0), which was developed with Snakemake v.7.32.4 and is available in Github (https://github.com/caceres-lab/GeONTIpe). Briefly, ONT reads with less than 5 kb of length and quality of 7 were filtered out using NanoFilt v.2.8.0 (De Coster and Rademakers 2023) and total sequence output, number of reads and N50 of each sequencing run were obtained with NanoPlot v.1.40.2 (De Coster and Rademakers 2023). Next, filtered reads were mapped against the hg38 reference genome with minimap2 v.2.23-r1111 (Li 2021) using the following parameters: ax map-ont -z 400,100 -r 100,1000 -- secondary=no. Unmapped and mapped reads with MAPQ < 20 were removed from the alignment file using samtools v.1.14 (Danecek et al. 2021), whereas reads mapping to each inversion region (external inversion breakpoint coordinates ± 100 kb) were selected. Specific inversion probes were then mapped to the reads of the corresponding region using Blastn v.2.12.0 with parameters -task megablast -outfmt "6 sstart send slen sstrand" -qcov_hsp_perc 90 -sorthits 4, and the inversion allele of the individual reads was determined based on the relative order, orientation and distance of the probes at both sides of the breakpoint. To ensure that genotypes for each inversion and sample are as reliable as possible, they were defined by the number of informative reads supporting each orientation as: (i) homozygote, if one orientation is supported by ≥5 reads for autosomes and Chr. X in females (corresponding to a binomial p-value of being heterozygote of less than 0.031), which is reduced to ≥2 reads for haploid Chr. X and Y in males, and there are <5% reads of the other orientation (to allow for the presence of 1-2 inconsistent minoritary reads in inversions with high coverage); (ii) heterozygote, when there are ≥1 O1 and O2 reads and they represent at least 15% of the total reads, to avoid incorrect genotype calls based in a few spurious reads; and (iii) low confident, when there are insufficient reads or a small proportion of reads (5-15%) of one orientation. Moreover, for low confident cases with <5 reads supporting the same inversion orientation, we analyzed known SNPs to test if both chromosomes are already represented in these few reads and recover additional genotypes. Biallelic SNPs within the inversion region were selected from dbSNP (Sherry et al. 2001) and those with <15% frequency or identified as homozygotes in the target samples in the 1KGP data were filtered out (Byrska-Bishop et al. 2022). SNPs present in informative reads with alleles concordant to those expected were used to calculate a distance between reads (value between 0 and 1) and create a dendrogram with the Hclust function of the R 4.1.2 Stats package. Read clusters showing a distance higher than 0.5 were considered as different haplotypes belonging to an homozygote for the supported orientation.

Additional SVs of more than 200 bp in the inversion regions were detected based on a >5% discrepancy between the expected and the observed distance among the normal combinations of the four probes (AB, CD, AC, BD, BC, and AD) in >15% of reads with the same orientation (Sup. Table 4). The distance difference was then used to classify the SV as either an insertion (Observed > Expected) or deletion (Observed < Expected) and to determine its size. The number of independent additional SVs was determined by removing the variants present in all O1 or O2 chromosomes (which represent possible errors in the reference genome or in the generated inverted reference) and duplicated SVs with the same size detected separately in O1 and O2 chromosomes in the same location (including insertions and deletions exchanging breakpoints between O1 and O2 chromosomes that can be moved by the inversion). Multiple insertions and deletions located in the same region with sizes varying by a similar amount were considered CNVs and counted as a single SV. Cases where no internal B and C probes were found and there was a size difference between A and D probes compatible with a deletion in >15 % of the reads were classified as AD deletions (complete deletion of inversion region) and one of the alleles of the individual was labelled as Del. Finally, when probes showed incompatible orientation conformations, the reads were classified as errors, suggesting the presence of potential new haplotypes that may require further analysis.

### Data analysis

Average sequencing enrichment in adaptive sampling was obtained by the ratio of the coverage of targeted and non-targeted regions in each run. In addition, to analyze the variation across different genomic regions with AS2 and AS3 in Monarch extracted DNA, we divided the genome in 10-kb windows classified based on their location inside or outside the targeted regions and calculated their enrichment by dividing their sequencing depth by the average depth of non-targeted windows in that run. Enriched non-targeted windows were identified as the top 1% average enrichment values across all AS2 or AS3 runs (which corresponds to an enrichment of 3.6 for AS2 and 3.8 for AS3), after excluding those within 100 kb of the targeted regions to account for the increased coverage due to reads extended from the targets. To assess the distribution of repetitive elements in these windows, the hg38 Segmental Dups and RepeatMasker annotations (including LINEs, SINEs, simple repeats, LTRs, satellites and low-complexity DNA) were obtained from the UCSC genome browser and the content of both types of elements in the windows was calculated separately. Windows were considered significantly enriched in SDs or repeats if the fraction of that type of repetitive element was higher than the 95% percentile of the genome distribution (corresponding to 53.5% for SDs or 89.3% for other repeats). Finally, we checked if the sequence identity between the enriched non-targeted windows and the AS3 target sequences was responsible for part of the unexpected enrichment by comparing them using Blast 2.12.0 (Altschul et al. 1990) with parameters -task blastn - qcov_hsp_perc 50 -perc_identity 85.

Comparison of ONT inversion genotypes with those of available predictions was done based on the inversions from the merged dataset detected in each of the previous studies depending on whether the information available included actual genotypes (Porubsky et al. 2022; Chaisson et al. 2019; Lerga-Jaso et al. in prep.) or just the presence or absence of the inversions (Levy-Sakin et al. 2019; Audano et al. 2019). Analysis of the Pangenome samples with available paths involved the conversion of the pggb draft (Liao et al. 2023) into sequences. Hg38 coordinates were used as input to select paths and nodes with Odgi v0.8.3 extract function (Guarracino et al. 2022) and odgi paths with parameter -f was applied to convert the odgi into FASTA format. The obtained output was used to run the inversion genotyping pipeline with default parameters, considering that there is only one sequence per haplotype.

## DATA ACCESS

All sequencing data generated in this study have been submitted to the NCBI Sequence Read Archive (SRA) (https://www.ncbi.nlm.nih.gov/sra) under accession numbers XXXX-XXXX. The code of the GeONTIpe inversion genotyping package is available in Github (https://github.com/caceres-lab/GeONTIpe). Inversion genotypes are available in the Supplemental Material and they have been deposited in dbVar (accession number XXXX).

## ACKNOWLEDGEMENTS

We thank Jim Thomas, Olga Francino, Michael Vieyra and the ONT Technical Support team and multiple ONT users for advice and technical support about DNA extraction and nanopore sequencing, Carles Acosta for help with the implementation of the genotyping pipeline in a supercomputer environment, David Porubsky and Jan Korbel for suggestions about the development of the Snakemake package, Odei Blanco for individual inversions quality control information, and Adrià Mompart for thorough GeONTIpe testing. The data processing and analysis tools have been developed, implemented and operated in collaboration with the Port d’Informació Científica (PIC) data center. PIC is maintained through a collaboration agreement between the Institut de Física d’Altes Energies (IFAE) and the Centro de Investigaciones Energéticas, Medioambientales y Tecnológicas (CIEMAT). This work was supported by research grants PID2019-107836RB-I00 and PID2022-137615OB-I00 to MC and PID2023-146193OB-I00 to MAS funded by the Agencia Estatal de Investigación of the Ministerio de Ciencia, Innovación y Universidades (MICIU/AEI/10.13039/501100011033, Spain) and the European Regional Development Fund (ERDF, EU), 2021 SGR 00526 support to research groups from the Departament de Recerca i Universitats (Generalitat de Catalunya, Spain) to MC, the predoctoral program AGAUR-FI Joan Oró grants 2021FI-B 00240 to RMP and 2023 FI-1 00746 to MDR funded by the Departament de Recerca i Universitats (Generalitat de Catalunya, Spain), as well as the European Social Plus Fund, a Maria Zambrano Grant from the MICIU (Spain) funded by the European Union-NextGenerationEU to KK, and Predoctoral Contract PRE-2020-092440 to IY (MICIU/AEI/10.13039/501100011033, Spain). MP is a Serra Húnter Fellow.

## AUTHOR CONTRIBUTIONS

MC conceived the project, devised the study and oversaw all the steps. RMP and IY developed the inversion genotyping pipeline with help from AS, JMU, MP and MC. RMP, MP, MDR and MC selected the inversion regions. KK, OC, MAS and MP performed the ONT sequencing. RMP, KK and MP analyzed the previously available and newly generated ONT sequencing data. RMP, KK, MP, and MC wrote the manuscript, and all the authors contributed comments to its final version.

